# Impacts of Self-Administered 3,4-Methylenedioxypyrovalerone (MDPV) Alone, and in Combination with Caffeine, on Recognition Memory and Striatal Monoamine Neurochemistry in Male Sprague-Dawley Rats: Comparisons with Methamphetamine and Cocaine

**DOI:** 10.1101/2024.01.31.578247

**Authors:** Robert W. Seaman, Kariann Lamon, Nicholas Whitton, Brian Latimer, Agnieszka Sulima, Kenner C. Rice, Kevin S. Murnane, Gregory T. Collins

**Affiliations:** Department of Pharmacology, The University of Texas Health Science Center at San Antonio, San Antonio, TX, United States; Department of Pharmacology, Toxicology and Neuroscience, Louisiana State University Health Sciences Center at Shreveport, Shreveport, LA, United States; Louisiana Addiction Research Center, Louisiana State University Health Sciences Center at Shreveport, Shreveport, LA, United States; Department of Psychiatry and Behavioral Medicine, Louisiana State University Health Sciences Center at Shreveport, Shreveport, LA, United States; Drug Design and Synthesis Section, Molecular Targets and Medications Discovery Branch, Intramural Research Program, National Institute on Drug Abuse and the National Institute on Alcohol Abuse and Alcoholism, National Institutes of Health, Bethesda, Maryland, United States; South Texas Veterans Health Care System, San Antonio, TX, United States

**Keywords:** MDPV, Bath Salts, Stimulants, Self-Administration, Recognition Memory, Neurochemistry

## Abstract

Recent data suggest that 3,4-methylenedioxypyrovalerone (MDPV) has neurotoxic effects; however, the cognitive and neurochemical consequences of MDPV self-administration remain largely unexplored. Furthermore, despite the fact that drug preparations that contain MDPV often also contain caffeine, little is known regarding the toxic effects produced by the co-use of these two stimulants. The current study investigated the degree to which self-administered MDPV, or a mixture of MDPV+caffeine can produce deficits in recognition memory and alter neurochemistry relative to prototypical stimulants. Male Sprague-Dawley rats were provided 90-min or 12-h access to MDPV, MDPV+caffeine, methamphetamine, cocaine, or saline for 6 weeks. Novel object recognition (NOR) memory was evaluated prior to any drug self-administration history and 3 weeks after the final self-administration session. Rats that had 12-h access to methamphetamine and those that had 90-min or 12-h access to MDPV+caffeine exhibited significant deficits in NOR, whereas no significant deficits were observed in rats that self-administered cocaine or MDPV. Striatal mono-amine levels were not systematically affected. These data demonstrate synergism between MDPV and caffeine with regard to producing recognition memory deficits and lethality, highlighting the importance of recapitulating the manner in which drugs are used (e.g., in mixtures containing multiple stimulants, binge-like patterns of intake).

## 1. Introduction

Stimulants remain one of the most popular classes of drugs used worldwide. Upon examination of the novel psychoactive stimulants identified in the drug supply, a large proportion of them are synthetic cathinones. Synthetic cathinones emerged on the il-licit drug market in the early 2000s in the form of “Bath Salts” preparations as popular “legal” alternatives to illicit psychostimulants, such as cocaine and methamphetamine, functioning in a similar manner as either inhibitors of monoamine reuptake or substrates for monoamine transporters [1–6]. Synthetic cathinones garnered much attention in the popular press due to a large number of adverse psychiatric (hallucinations, aggression, paranoia) and physiological (tachycardia, hyperthermia) effects as-sociated with the use of various “Bath Salts” preparations [7–10]. One of the most commonly reported synthetic cathinones found in “Bath Salts” preparations is 3,4-methylenedioxypyrovalerone, or MDPV, a monoamine reuptake inhibitor that is ∼750 times more potent at inhibiting reuptake at the dopamine transporter than the serotonin transporter [10–13]. Consistent with reports of MDPV being widely used, it has been demonstrated to share discriminative stimulus properties with other commonly used stimulants (e.g., cocaine, methamphetamine) [14–19], and readily maintains self-administration, acting as a highly effective reinforcer in both rodents and non-human primates [11,20–25].

Whether the chronic use of synthetic cathinones could produce long-term toxicities, as is observed following the chronic use of other stimulants, is unclear. In vitro data suggest that MDPV can produce oxidative stress and mitochondrial dysfunction ultimately leading to apoptosis in SH-SY5Y cells [26–30]. Additionally, MDPV has been shown to produce cytotoxicity in bovine brain microvessel endothelial cells, suggesting that MDPV could produce deficits in blood-brain barrier integrity and/or function [31]. To date, the vast majority of studies investigating adverse effects of MDPV have employed either relatively short daily access sessions (e.g., 90-min) or non-contingent exposure. Though deficits in novel object recognition have been reported following a “binge” regimen of three non-contingent injections of 1 mg/kg MDPV per day, for ten days [32], case reports suggest that “Bath Salts” users tend to engage in repeated bouts of drug-taking behavior that can last days or even weeks [33–37]. Accordingly, incorporating longer access sessions (e.g., 12-h/day) to response-contingent MDPV might better model patterns of MDPV use reported by humans and aid in detailing possible long-term consequences of MDPV use. A study that employed a more translationally relevant model of MDPV exposure showed that intravenous self-administration of MDPV over 5 96-h sessions was able to both disrupt recognition memory and increase markers of neurodegeneration in both the perirhinal and entorhinal cortices, relative to rats that only had access to saline infusions [38]. However, it remains unclear how those deficits produced by MDPV would compare to those in rats that had extended access to other stimulants (e.g., methamphetamine or cocaine), or if MDPV was ad-ministered in combination with other common “Bath Salts” constituents, such as caffeine. The latter is particularly important given both the frequency with which MDPV is mixed with caffeine [10,12], as well as a number of studies that suggest caffeine is able to synergistically enhance the reinforcing, relapse-related, and discriminative stimulus effects of MDPV in rats [14,39,40].

The current studies begin to address these gaps in knowledge by integrating a 12-h access self-administration procedure with the novel object recognition assay and ex vivo neurochemical analyses in order to address the following hypotheses: (1) 12-h access to MDPV and MDPV+caffeine will result in escalations of intake akin to those observed in rats self-administering methamphetamine and cocaine; (2) 12-h access to methamphetamine, MDPV, and MDPV+caffeine will result in deficits in recognition memory; (3) 12-h access to methamphetamine, MDPV, and MDPV+caffeine will deplete striatal levels of dopamine; and (4) the addition of caffeine to MDPV will produce greater deficits in recognition memory and striatal dopamine neurochemistry than are observed in rats self-administering MDPV alone.

## 2. Materials and Methods

### Self-administration

#### Subjects

84 male Sprague-Dawley rats (275–300 g upon arrival) were purchased from Envigo (Indianapolis, IN, United States) and maintained in a temperature- and humidity-controlled room. Rats were individually housed and maintained on a 14/10 h light/dark cycle. All experiments were conducted at approximately the same time each day. Rats were provided ad libitum access to Purina rat chow and water. All studies were carried out in accordance with the Institutional Animal Care and Use Committees of the University of Texas Health Science Center at San Antonio and the eighth edition of the Guide for Care and Use of Laboratory Animals (National Research Council (United States) Committee for the Update of the Guide for the Care and Use of Laboratory Animals 2011).

#### Surgery

Rats were anesthetized with 2–3% isoflurane and prepared with chronic indwelling catheters in the left femoral vein, as previously described [41,42]. Catheters were tunneled under the skin and attached to a vascular access button placed in the mid-scapular region. Immediately following surgery, rats were administered Excede (20 mg/kg; SC) to prevent infection. Rats were allowed 5–7 days to recover during which time catheters were flushed daily with 0.5 ml of heparinized saline (100 U/ml). Thereafter, catheters were flushed daily with 0.2 ml of saline prior to and 0.5 ml of heparinized saline after the completion of self-administration sessions.

#### Apparatus

All self-administration experiments were conducted in standard operant conditioning chambers located within ventilated, sound-attenuating enclosures (Med Associates, Inc., St. Albans, VT). Each chamber was equipped with two response levers located 6.8 cm above the grid floor and 1.3 cm from the right or left wall. Visual stimuli were pro-vided by two sets of green, yellow, and red LEDs, one set located above each of the two levers, and a white house light located at the top center of the opposite wall. Drug solutions were delivered by a variable-speed syringe pump through Tygon tubing connected to a stainless-steel fluid swivel and spring tether, which was held in place by a counterbalanced arm. Experimental events were controlled, and data were collected using MED-PC IV software and a PC-compatible interface (Med Associates, Inc.).

### Self-administration

Rats were initially allowed to respond for the presentation of grain pellets (45 mg) un-der a fixed ratio (FR) 1 schedule of reinforcement during ten, daily, 90-min sessions. Illumination of green and red LEDs above the active lever (left or right; counterbalanced across rats) signaled grain pellet availability and completion of the response requirement resulted in the delivery of a grain pellet, coinciding with the illumination of the houselight as well as the yellow, green, and red LEDs above the active lever, and initiation of a 5-s timeout throughout which responding had no scheduled consequences. Upon meeting acquisition criteria (≥15 grain pellets delivered for two consecutive days with ≥80% responding occurring on the active lever versus the inactive lever), a yellow LED was illuminated above the previously inactive lever signaling availability of an infusion of either saline, cocaine (0.32 mg/kg/infusion), methamphetamine (0.1 mg/kg/infusion), MDPV (0.032 mg/kg/infusion), or a mixture of MDPV+caffeine (0.032 mg/kg/infusion and 0.11 mg/kg/infusion, respectively), under an FR 1 schedule of reinforcement during at least 10, daily, 90-min sessions. Completion of the response requirement resulted in a drug infusion (0.1 ml/kg over ∼1 s), illumination of the houselight as well as the yellow, green, and red LEDs above the active lever, and initiation of a 5-s timeout throughout which drug was made unavailable. Throughout the remainder of the study, responding on the lever signaled by green and red LEDs resulted in the delivery of a grain pellet, and responding on the alternate lever signaled by a yellow LED resulted in an infusion of drug (or saline). Response requirements for both grain pellets and drug (or saline) were subsequently increased to an FR 5 schedule where they remained for the duration of the study. Six days after rats were maintained under an FR 5 schedule of reinforcement, the session duration for half of the rats from the saline, cocaine, methamphetamine, MDPV, and MDPV+caffeine groups was increased from 90 min to 12 h; for the other rats, session durations remained at 90 min throughout. Self-administration sessions were conduct-ed five days per week, for six weeks.

### Drugs

Racemic MDPV was synthesized by Agnieszka Sulima and Kenner Rice (Bethesda, MD). Cocaine was provided by the National Institute on Drug Abuse Drug Supply Program. D-methamphetamine and caffeine were purchased from Sigma-Aldrich (St. Louis, MO, United States). All drugs were dissolved in physiological saline.

### Novel Object Recognition

Novel object recognition (NOR) was conducted in custom-built arenas with black walls and floors (43 cm x 56 cm x 38 cm; W x L x H). Webcams were situated above the arenas to record sessions for scoring at a later time. Rats underwent habituation, training, and test phases. During habituation, rats were allowed to freely explore an empty arena for 30 min (no objects were placed in the arena). Twenty-four hours later, a training session was conducted in which rats were placed into an arena with two identical objects placed on opposite sides of the arena, 20 cm apart, and allowed to explore the objects for 5 min before being returned to their home cages. The testing phase occurred 60 min later wherein one of the two identical objects (counterbalanced for left and right sides) was replaced by a different object (novel object), after which rats were allowed to explore both objects for 3 min. The objects used were a metallic can, an aluminum foil-wrapped cardboard cone, a green striped aluminum foil-wrapped cardboard cone, a plastic bottle, a miniature wooden birdhouse, and a miniature plastic sandcastle. Pi-lot studies suggested that rats had no preference for any single object. The objects acting as familiar or novel were pseudorandomized. Exploration was defined as directing the nose toward the object at a distance of less than 2 cm [43]. The arena and the objects were cleaned between each trial with an alcohol-based cleaner. Recognition memory was quantified with a NOR score ([time spent with the novel object - time spent with the familiar object] / total exploration time). NOR was conducted prior to any drug self-administration and then three weeks following the final self-administration session. Throughout the 3-week drug-free period, rats remained in their home cages and were flushed daily.

### Neurochemistry

To quantify monoamine neurochemistry in the striatum, rats were euthanized via rapid decapitation, and brains were removed. Brains were immediately frozen on dry ice and stored at −80ᵒC for subsequent analyses. Brains were thawed at 4ᵒC, placed in an ice-cold rat brain matrix, and sliced into coronal sections (1 mm thick). These slices were next placed flat on a cold plate over ice. Using a 1.5 mm diameter tissue biopsy-punch, regions of interest were taken from individual slices, as described previously [44].

Frozen tissues were weighed, sonically disrupted in 100 µl of 0.3 N HClO4, and centrifuged for 10 minutes at 4ᵒC. A 100 µl aliquot of the supernatant was placed in a Waters 717plus autosampler maintained at 10ᵒC, and 50µl was injected onto a Waters Spherisorb ODS 5µm column, 4.6 x 250mm (28ᵒC), 0.1M sodium phosphate with 0.1mM EDTA, 0.3mM OSA, and 3.75% acetonitrile (pH adjusted to 3.05 with O-Phosphoric acid) running at a flow rate of 0.8 mL/min. Amperometric detection was accomplished with a BAS LC-4C detector equipped with a glassy carbon electrode (set to 780mV) and the signal was analyzed on Waters Empower software (ver. 2). Absolute tissue concentrations (ng/mg) for the monoamine neurotransmitters dopamine, 3,4-dihydroxyphenylacetic acid (DOPAC), homovanillic acid (HVA), serotonin, 5-hydroxyindoleacetic acid (5-HIAA), norepinephrine, and epinephrine were deter-mined by comparison with external standard curves and corrected for tissue weight, as described previously [44].

### Data Analyses

Weekly intake data represent the mean (±S.E.M) level of drug (mg/kg) or food (pellets) earned across the 5 daily sessions. Intake data are plotted as a function of study week and were analyzed via two-way (access condition x experimental week) repeated measure analysis of variance (ANOVA) and post-hoc Holm-Sidak test. Daily intake data are presented as the mean ±S.E.M. Total experimental intake data are presented for individual subjects and were analyzed via Student’s t-tests (90-min versus 12-h access). Novel object recognition scores, total exploration time, and striatal, dopamine neurochemistry data are shown for individual subjects and were analyzed via two-way (access condition x experimental week) ANOVA and post-hoc Holm-Sidak test. Given the relatively small quantities of norepinephrine and serotonin in the striatum, and their respective metabolites, these neurochemistry data are described and shown but not analyzed statistically.

## 3. Results

### Self-administration

The average daily intakes for each reinforcer during 90-min or 12-h access sessions across each of the 6 experimental weeks are presented in Figure 1. Overall, there were main effects of session duration, with rats provided 12-h access to drug achieving greater average levels of drug intake per session relative to rats provided 90-min access, revealed by repeated-measures ANOVAs with regard to cocaine (F(1,15)=22.98, p<0.05; upper left), methamphetamine (F(1,16)=17.62, p<0.05; upper right), MDPV in rats that had access to MDPV alone (F(1,15)=11.56, p<0.05; middle left), and MDPV in rats that had access to a mixture of MDPV+caffeine (F(3,26)=18.1, p<0.05; middle right), caffeine in rats that had access to a mixture of MDPV+caffeine (F(3,26)=18.1, p<0.05; bottom left), and grain pellets (F(1,15)=11.56, p<0.05; bottom right).

**Figure 1.**
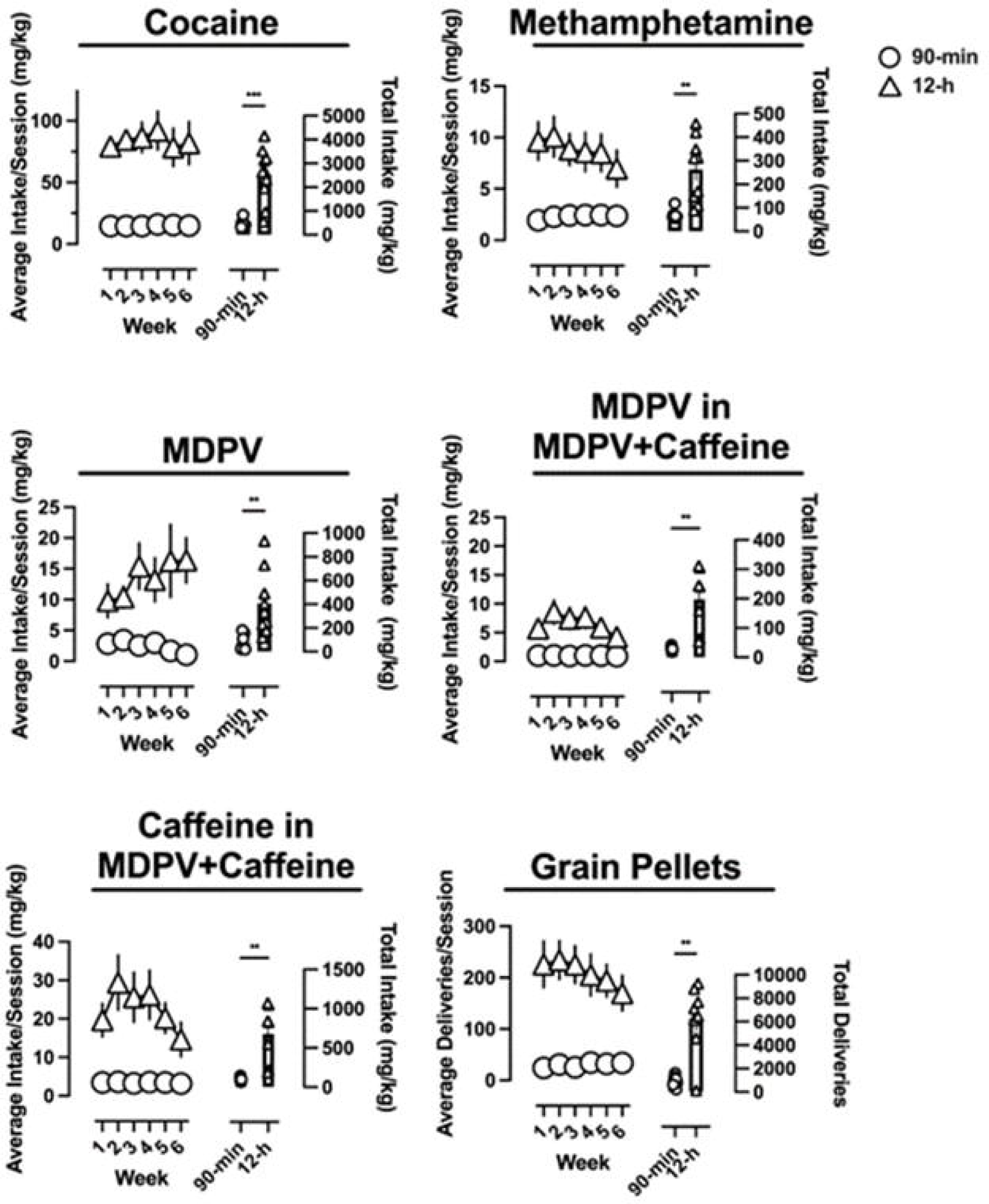
Average drug intake (or number of pellet deliveries for rats that had access to saline and grain pellets) (left ordinate) per session as a function of experimental week. Total experimental in-take in rats that had access to each reinforcer for 90-min or 12-h is represented on the right ordinate. Triangles represent rats that had 12-h access, and circles represent rats that had access for 90-min. Data represent the mean ±S.E.M. ** denotes p<0.01, *** denotes p<0.001

When evaluated based on average intake per session as a function of week, cocaine self-administration remained stable over the course of the experiment, regardless of access condition, with no main effect of experimental week (F(5,75)=0.47, p>0.05). Although there was no main effect of experimental week (F(5,80)=0.81, p>0.05) aver-age methamphetamine intake per session decreased over the course of the study in rats provided 12-h access, but slightly increased in rats provided 90-min access.

The opposite was true for MDPV, in that, rats provided 12-h access increased their average session intake over the course of the study, whereas rats provided 90-min access slightly decreased their MDPV intake, with no main effect of experimental week observed (F(5,75)=0.67, p>0.05). Rats responding for MDPV+caffeine earned fewer over-all infusions than rats self-administering MDPV alone, meaning less MDPV intake. In-take of both MDPV and caffeine in rats self-administering a mixture of both drugs resulted in an initial increase in drug intake followed by a decrease over the course of the experiment. There was a significant effect of experimental week for intake of MDPV (F(5,65)=3.1, p<0.05) as well as of caffeine (F(5,65)=3.1, p<0.05). Post-hoc tests revealed significant increases in average MDPV and caffeine intake per session between weeks 1 and 2 (p<0.05). 12-h access to grain pellet delivery resulted in an overall decrease over the course of the experiment although no significant effect of experimental week (F(5,75)=1.6, p>0.05).

Perhaps unsurprisingly, total drug intake (i.e., over the course of the entire study) was significantly greater in rats having 12-h access to cocaine (t=5.7, p<0.05), methamphetamine (t=4.2, p<0.05), MDPV (alone) (t=3.6, p<0.05), MDPV (MDPV+caffeine) (t=4.1, p<0.05), caffeine (MDPV+caffeine) (t=4.1, p<0.05), and grain pellets (t=5.2, p<0.05) relative to rats that were provided 90-min access.

To examine how drug-taking patterns change from day-to-day, and whether these patterns differ depending on the drug(s) being self-administered, daily intake data are shown in Figure 2. Cocaine self-administration minimally fluctuates across days over the course of the study, regardless of access condition. Rats self-administering methamphetamine, however, self-administered more drug earlier in the experimental week, and then decrease drug intake over the course of the week.

**Figure 2.**
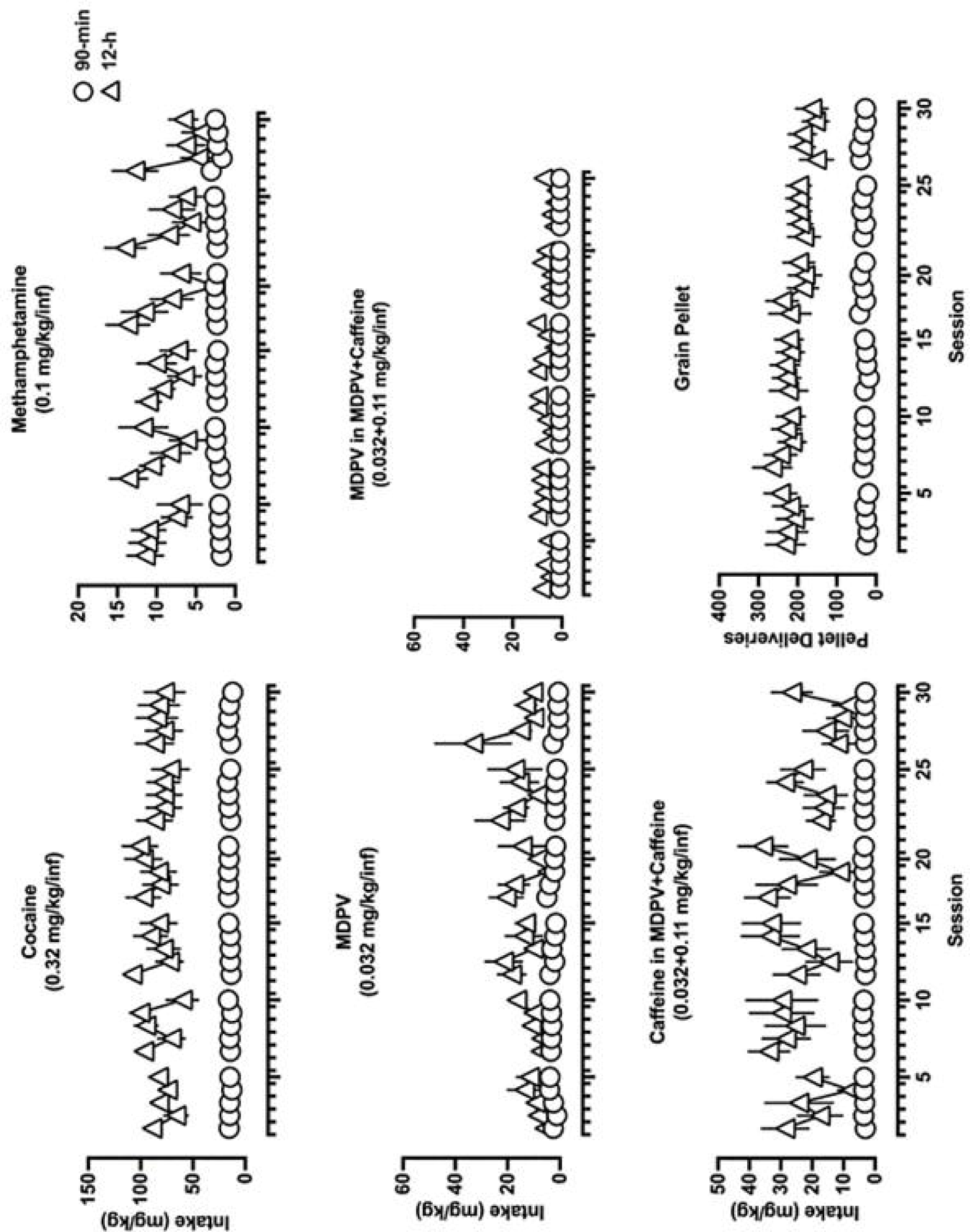
Daily drug intake (or number of pellet deliveries for rats that had access to saline and grain pellets) over the course of the experiment. Triangles represent rats that had 12-h access and circles represent rats that had 90-min access. Data represent the mean ±S.E.M.

Although this is more apparent in rats provided 12-h access to methamphetamine, a similar pattern can be observed, particularly in week 6, in rats provided only 90-min access to methamphetamine. 12-h access to MDPV initially resulted in rats increasing daily intake for the first two weeks, and then transitioned to a pattern of in-take not dissimilar from methamphetamine (i.e., more drug intake earlier in the week than towards the end). Similarly, by week three, rats self-administering MDPV in 90-min sessions began self-administering more drug early in the week and then reducing intake over the course of the week. 12-h access to MDPV+caffeine resulted in far less MDPV intake, but patterns of responding were similar. Rats self-administered more drug early in the week, followed by a subsequent decrease, and then often an increase by the end of the week (i.e., a “V” shaped pattern of intake within an experimental week. Rats responding for grain pellets exhibited relatively stable levels of responding across days over the course of the study, regardless of access condition.

### Novel Object Recognition

Recognition memory was probed at baseline (i.e., before any drug self-administration) and following a 3-week drug-free period. Two-way repeated measures ANOVAs revealed significant main effects of experimental week in rats that had self-administered methamphetamine (F(1,16)=7.2, p<0.05) (Figure 3; upper right) or a mixture of MDPV+caffeine (F(1,25)=10.4, p<0.05) (Figure 3; middle right) with post-hoc tests revealing a significant decrease in NOR score from baseline in rats that had 12-h access to methamphetamine, and rats that had either 90-min or 12-h access to MDPV+caffeine. In contrast, no main effects of experimental week on NOR score were observed in rats that had access to cocaine (F(1,17)=0.30, p>0.05) (Figure 3; upper left), grain pellets (F(1,14)=0.14, p>0.05) (Figure 3; bottom left), or MDPV (F(1,15)=0.3, p>0.05) (Figure 3; middle left).

**Figure 3.**
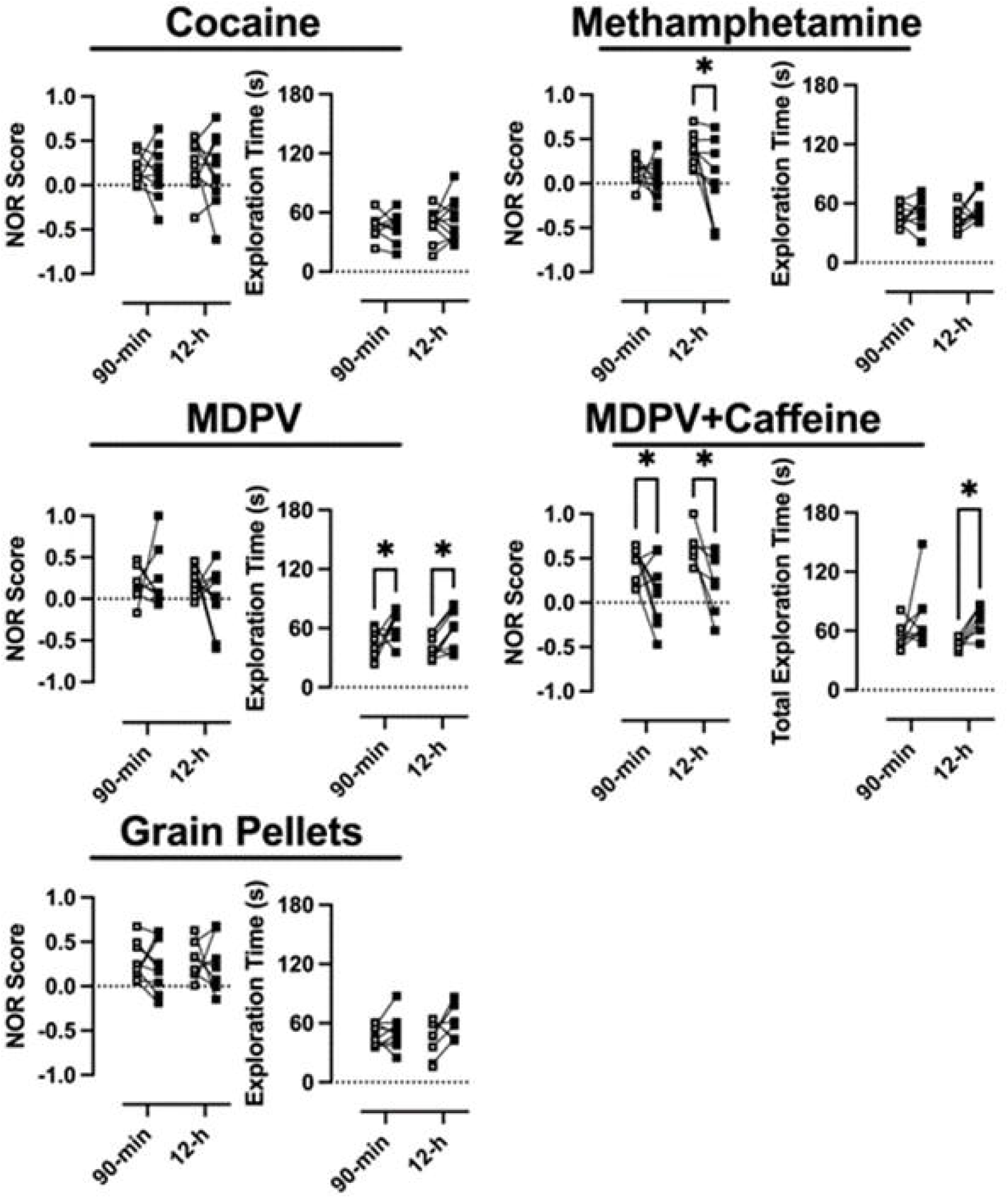
Average NOR score and total exploration time at baseline (prior to any drug exposure) (open squares) versus following a three-week drug-free period (closed squares). Data represent the mean ±S.E.M. * denotes p<0.05

Significant main effects of experimental week with regard to total exploration time were observed in rats that had self-administered MDPV (F(1,15)=13.1, p<0.05) or MDPV+caffeine (F(1,25)=9.5, p<0.05), with post-hoc tests revealing significant increases in total exploration time (p<0.05) in rats that had either 90-min or 12-h access to MDPV, or 12-h access to MDPV+caffeine.

### Striatal Monoamine Neurochemistry

#### Dopamine

Long-term effects of 90-min or 12-h access to each reinforcer on striatal dopamine neurochemistry were assessed following a 3-week drug-free period (Figure 4). There was a significant main effect of drug on levels of dopamine (F(4,77)=2.6, p<0.05) with post-hoc tests revealing a significant decrease in dopamine in rats allowed 90-min access to cocaine, relative to those that had 90-min access to grain pellets (Figure 4; top). No other significant changes in dopamine levels were observed as a function of self-administration of any other drug. With regard to the dopamine metabolite, 3,4-dihydroxyphenylacetic acid (DOPAC), there was a significant main effect of access condition (F(1,77)=7.4, p<0.05) and an interaction between access condition and drug (F(4,77)=2.9, p<0.05), however, post-hoc tests revealed no significant differences in DOPAC levels in the striatum of rats that had either 90-min or 12-h access to any drug versus those having access to grain pellets (Figure 4; upper middle). A two-way ANOVA also revealed significant main effects of both access condition (F(1,73)=84, p<0.05) and drug (F(4,73)=15.1, p<0.05), as well as an interaction (F(4,73)=13.9, p<0.05) on striatal levels of homovanillic acid (HVA) (Figure 4; lower middle). Post-hoc analyses revealed significant decreases in HVA following 90-min access to cocaine, MDPV, or MDPV+caffeine, whereas 90-min access to methamphetamine resulted in a significant increase in HVA, relative to levels in rats responding for grain pellets. There were no significant differences in HVA levels following 12-h access to any drug, relative to grain pellets. Finally, with regard to dopamine turnover (i.e., levels of DOPAC / levels of dopamine), there were significant main effects of both access condition (F(1,77)=17.8, p<0.05) and drug (F(4,77)=6.1, p<0.05), as well as an interaction (F(4,77)=2.9, p<0.05) (Figure 4; bottom). Post-hoc analyses revealed a significant in-crease in dopamine turnover in rats allowed 90-min access to methamphetamine, relative to those responding for grain pellets.

**Figure 4.**
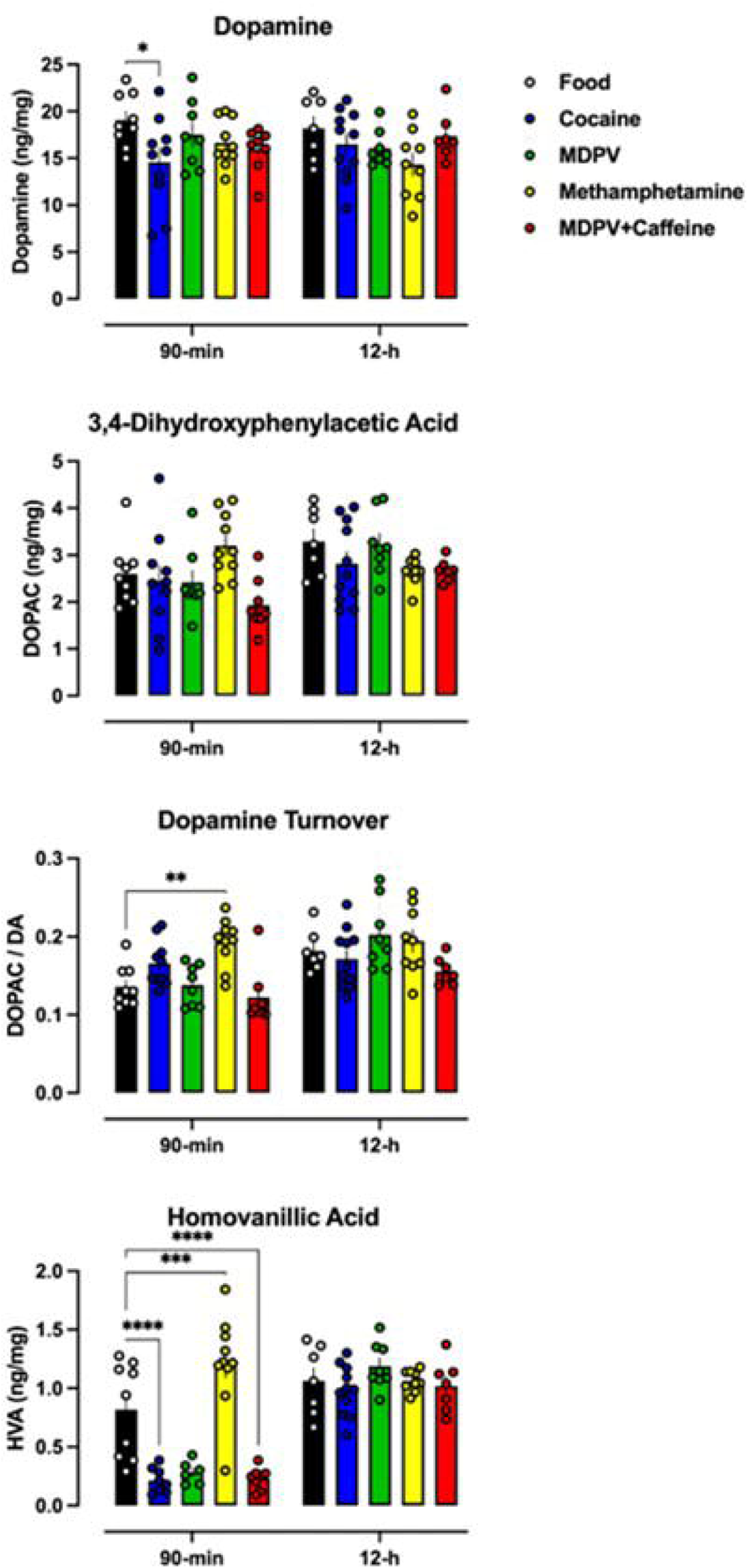
Average levels of dopamine and dopamine metabolites as a function of both reinforcer and access duration. Data represent the mean ±S.E.M. * denotes p<0.05, ** denotes p<0.01, *** denotes p<0.001, **** denotes p<0.0001

#### Norepinephrine

Apparent increases in striatal norepinephrine levels were observed in rats allowed 90-min or 12-h access to methamphetamine, relative to those that had 90-min access to grain pellets (Figure 5; top). With regard to striatal levels of epinephrine, seemingly small reductions were observed in rats that had 90-min access to any of the drugs, as compared to rats that had 90-min access to grain pellets. In contrast, only 12-h access to methamphetamine and MDPV+caffeine resulted in apparent decreases in striatal epinephrine (Figure 5; middle). With regard to norepinephrine turnover (i.e., levels of epinephrine / levels of norepinephrine), there were apparent increases in norepinephrine turnover in rats allowed 90-min access to any of the drugs, relative to those that had access to grain pellets. In contrast, 12-h access to any of the drugs did not appear to produce large differences in norepinephrine turnover compared to what was observed in rats responding for grain pellets in 12-h sessions (Figure 5; bottom).

**Figure 5.**
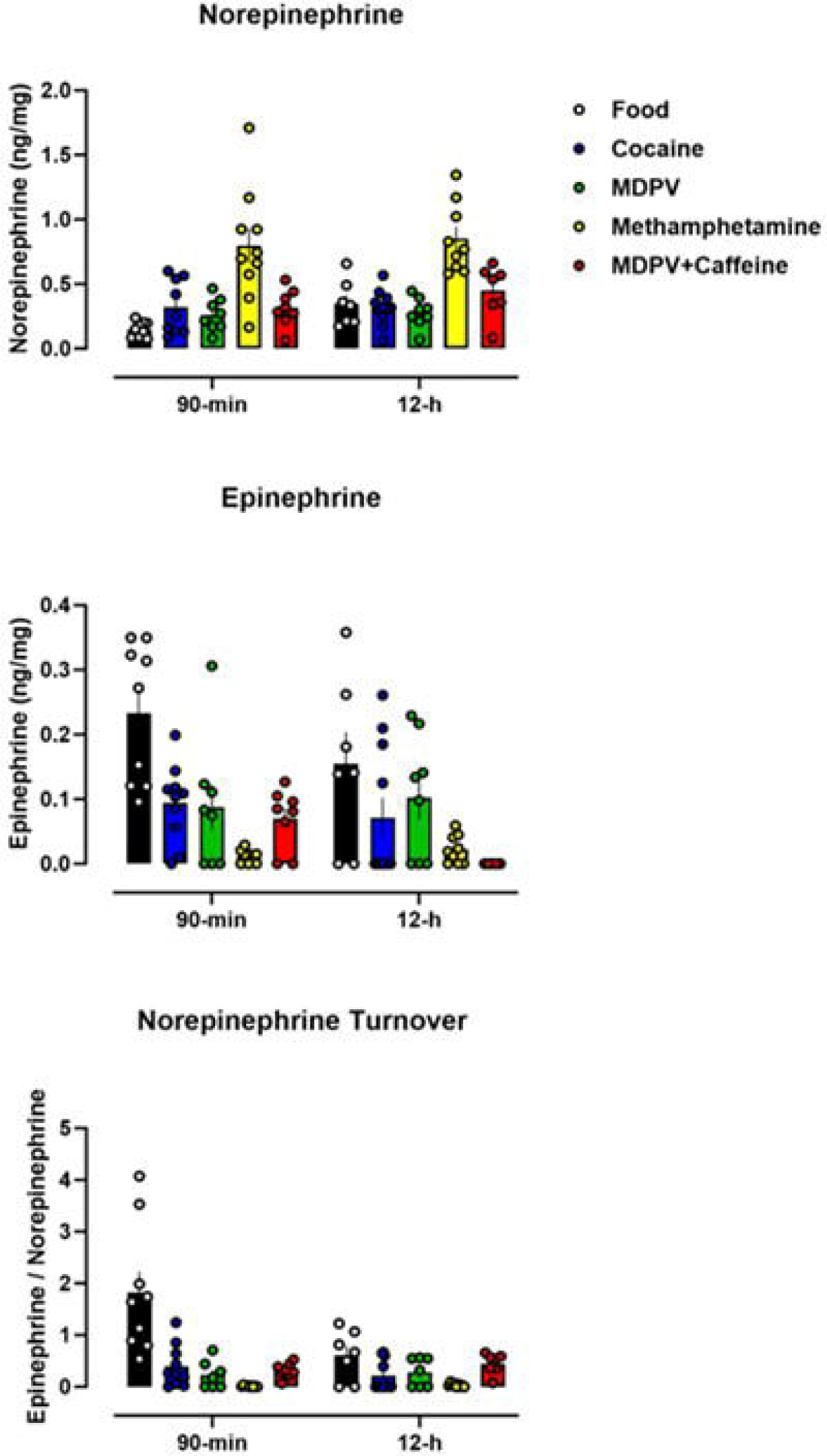
Average levels of norepinephrine and norepinephrine metabolites as a function of both reinforcer and access duration. Data represent the mean ±S.E.M.

#### Serotonin

When levels of striatal serotonin were assessed, there was an apparent reduction in serotonin observed for rats allowed 90-min or 12-h access to any of the drugs, relative to those that had access to grain pellets (Figure 6; top). Unlike what was observed with serotonin, none of the drug access conditions appeared to affect the striatal levels of the serotonin metabolite, 5-hydroxyindoleacetic acid (5-HIAA) (Figure 6; middle). With regard to serotonin turnover (i.e., levels of 5-HIAA / levels of serotonin), slight increases were observed for rats allowed 12-h access to any of the drugs, relative to those responding for grain pellets. In contrast, when levels of striatal serotonin turnover were compared for rats from the 90-min access conditions, there were no apparent differences, regardless of reinforcer (Figure 6; bottom).

**Figure 6.**
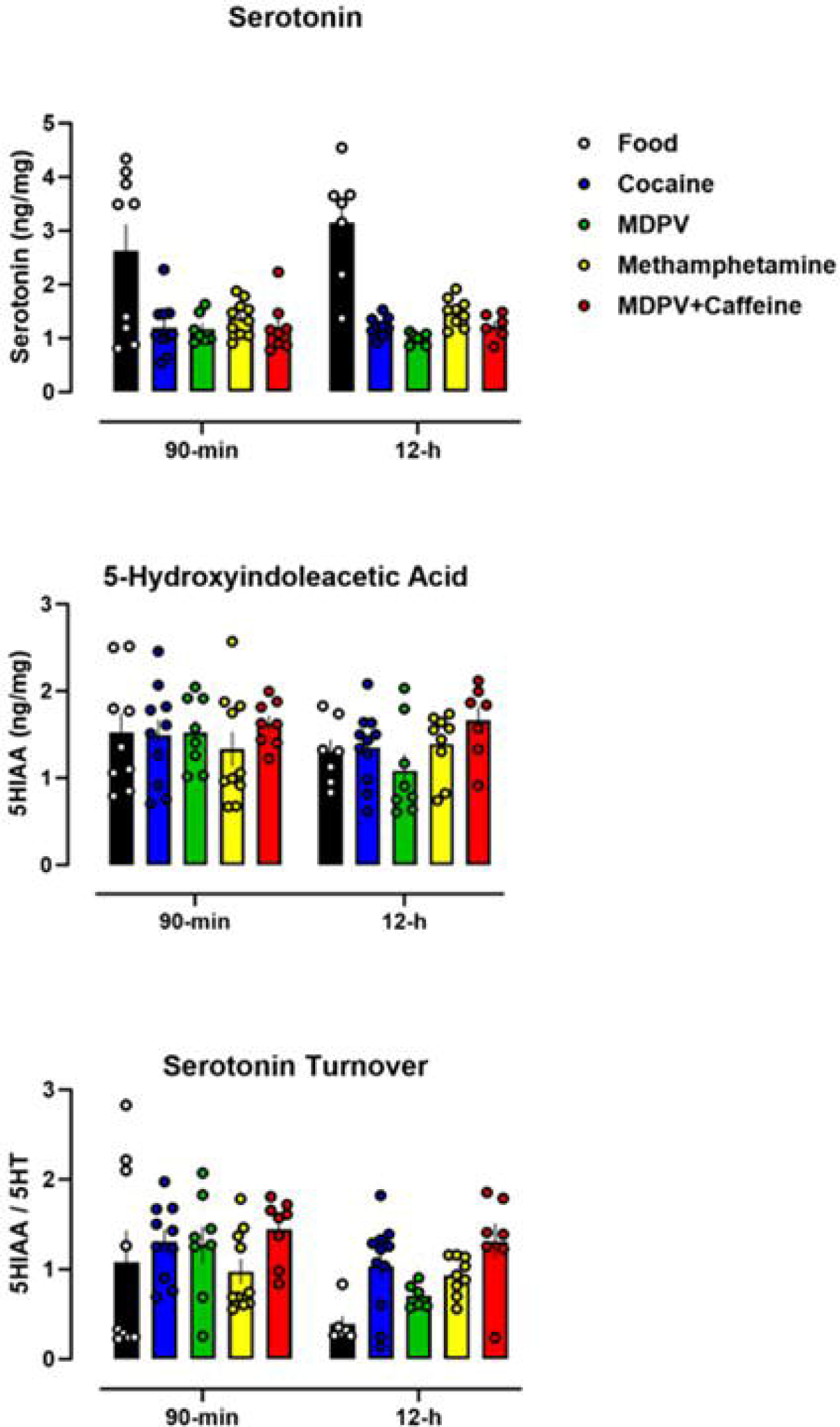
Average levels of serotonin and serotonin metabolites as a function of both reinforcer and access duration. Data represent the mean ±S.E.M.

## 4. Discussion

Although MDPV has been reported to be one of the most prominent synthetic cathinones in “Bath Salts” preparations [10,12,13], relatively little is known regarding consequences of its use. The current studies utilized an uncapped, 12-h access self-administration paradigm to compare patterns of drug intake, changes in recognition memory, and alterations in striatal monoamine neurochemistry in rats self-administering MDPV, cocaine, or methamphetamine. In addition, because “Bath Salts” preparations often contain other psychoactive substances, such as caffeine, these studies also included a mixture of MDPV+caffeine in order to determine if the presence of caffeine altered the behavioral and neurochemical consequences of MDPV self-administration. There were five main findings: 1) Patterns of self-administration were drug-dependent, with cocaine maintaining relatively stable levels of responding across an experimental week, whereas rats self-administering methamphetamine, MDPV, and MDPV+caffeine exhibited “binge-like” patterns of drug intake, self-administering larger amounts of drug earlier in the week versus later in the week; 2) the addition of caffeine to MDPV resulted in less MDPV intake than rats provided access to the same dose of MDPV in isolation; 3) 12-h access to methamphetamine resulted in deficits in recognition memory not observed in rats provided 12-h access to cocaine or MDPV; 4) 90-min and 12-h access to a mixture of MDPV and caffeine resulted in deficits in recognition memory, despite less overall MDPV intake; and 5) overall, striatal monoamine neurochemistry was not systematically altered as a function of access condition, drug being self-administered, or levels of intake. Taken together, these data suggest that extended access to MDPV alone is insufficient to pro-duce long-lasting deficits in cognition or changes in monoamine neurochemistry in the striatum, whereas the addition of caffeine to self-administered MDPV can result in cognitive deficits, even in rats allowed only 90-min access per day, highlighting the importance of evaluating drug mixtures rather than just single drugs, in isolation, for drugs that are commonly co-used.

In an attempt to recapitulate the “binge-and-crash” patterns often reported in human stimulant users, the current studies allowed rats uncapped access to cocaine, methamphetamine, MDPV, or MDPV+caffeine, during either 90-min or 12-h sessions con-ducted five days per week, for six weeks. Regarding patterns of intake across a given experimental week, cocaine intake tended to be relatively stable across the course of the experiment, with similar levels of intake occurring each session. This was in contrast to what was observed in rats self-administering methamphetamine, MDPV, or MDPV+caffeine in 12-h sessions, wherein intake was unevenly distributed throughout a given experimental week (e.g., greater levels of drug intake early in the week followed by days of far less drug intake), possibly recapitulating some aspects of stimulant use patterns in humans. Of note, rats allowed access to a mixture of MDPV+caffeine self-administered less MDPV than rats allowed access to MDPV alone. Although the drivers of this pattern of intake, and why it seemingly did not develop in rats self-administering cocaine, are outside of the scope of this study, it is likely the culmination of large levels of drug intake that disrupted a plethora of physiological (e.g., circadian, feeding, cardiovascular) functions. Perhaps during sessions of little to no drug intake, these physiological functions were restored sufficiently that drug in-take would be greater the following day. This pattern of intake largely recapitulates the “binge-and-crash” patterns of stimulant use reported in human users. As mentioned previously, few studies have allowed rats greater than 6-h access to methamphetamine, and those that have typically either cap the number of infusions that can be earned in a single session, or integrate periods of drug unavailability (e.g., 12-h access every other day, 1-h drug-free period every 3-h), perhaps reducing the likelihood of such patterns of intake developing.

One study that has evaluated extended access to MDPV self-administration, wherein rats were allowed to self-administer MDPV within five 96-h sessions each separated by a three-day drug-free period found that MDPV intake was demonstrated to significantly increase over time relative to rats that had access to infusions of saline. However, it should be noted that rats had not previously acquired self-administration behavior prior to starting the first 96-h self-administration session that intake in subsequent sessions was compared to, likely contributing to the “escalation of intake” that was observed [38]. Those data were presented as intake per 96-h session, so the resolution with which one can compare those data with the current data with regard to patterns of intake across 12-h is limited. In the current studies, self-administration of a mixture of MDPV+caffeine (0.032 mg/kg/inf+0.11 mg/kg/inf, respectively) resulted in fewer in-fusions being earned and therefore less MDPV intake than self-administration of MDPV (0.032 mg/kg/inf) alone. This is likely due to MDPV and caffeine, at these doses, having an additive effect with regard to reinforcing effects [40] (i.e., the mixture of MDPV+caffeine acted as a functionally larger reinforcer, maintaining fewer infusions under a fixed ratio schedule of reinforcement, akin to what would be observed if a larger MDPV dose was made available) or that the amount of caffeine being self-administered had rate-decreasing effects. It should be noted that 12-h access to MDPV+caffeine proved to be lethal in two rats, whereas no other condition in the cur-rent studies resulted in lethality. In these two subjects, total caffeine intake averaged ∼100 mg/kg in 12-h for multiple sessions before overdosing. Lethality resulting from that level is consistent with the LD50 reported for intravenous caffeine, 105 mg/kg [45]. Our laboratory and others have demonstrated that intravenous caffeine, when self-administered in isolation, does not maintain intake levels that are consistent with a caffeine overdose, suggesting that the addition of MDPV aided in increasing caffeine intake to the point of reaching potentially lethal levels [46–49]. In addition, MDPV and caffeine could interact synergistically with regard to toxic effects, decreasing the dose of either drug necessary to be lethal. Nevertheless, given that few studies have pro-vided rats with extended access to caffeine alone, and the current studies only evaluated one mixture of MDPV+caffeine, future work should better characterize toxic interactions between MDPV and caffeine using extended-access self-administration as an exposure paradigm.

Because long-term use of stimulants has been linked to cognitive deficits across a number of domains, the current studies sought to directly compare the impact of self-administered methamphetamine, cocaine, MDPV, and MDPV+caffeine on cognition. Though these effects tend to be more prominent with methamphetamine than cocaine, preclinical data suggest the use of MDPV can result in cognitive deficits. Thus, to examine the lasting consequences of various stimulant self-administration histories on cognitive performance related to recognition memory, novel object recognition was evaluated three weeks following the final self-administration session. Consistent with previous reports, 90-min or 12-h access to cocaine did not produce deficits in recognition memory, whereas 12-h (but not 90-min) access to methamphetamine did [50–54] (but see [55]). This is consistent with reports that drugs acting as reuptake inhibitors produce fewer cognitive deficits relative to those acting as substrates for monoamine transporters. Similar to what was observed with cocaine, and consistent with its mechanism of action as a monoamine reuptake inhibitor, self-administered MDPV alone (in either 90-min or 12-h sessions) did not produce deficits in recognition memory relative to the pre-drug baseline. This is in contrast to a previous report in rats wherein blunted recognition memory was observed following multiple 96-h self-administration sessions wherein MDPV was available [38]. This discrepancy could be the result of differences in drug availability (96-h sessions versus 12-h sessions), using rats that had access to saline as a comparator rather than rats that responded for grain pellets (i.e., controlling for a history of operant behavior), or in the time allowed to explore the two identical objects during the training phase of the novel object recognition assay prior to the testing phase (3-min versus 5-min). Previous work suggests that the lack of an effect of 12-h access to either reuptake inhibitor on recognition memory could be due to the task not requiring enough of a cognitive burden to unveil deficits in memory. Indeed, it has been shown that 6-h access to cocaine produced deficits in recognition memory when the interval between the training and test phases was 24-h, but not when the interval between training and test phases was only 3-h [56,57]. Future work should use a battery of cognitive assays to attain better resolution in characterizing functional cognitive deficits following stimulant self-administration.

Interestingly, a history of either 90-min or 12-h self-administration of a mixture of MDPV+caffeine did result in deficits in recognition memory, relative to baseline, with the caveat that the average pre-drug baseline NOR score in these subjects was larger than that in other groups. Although identifying the underlying mechanisms contributing to synergism between MDPV and caffeine with regard to persistent effects on memory is outside the scope of these studies, the idea that synergism between MDPV and caffeine exists with regard to producing memory deficits is consistent with previous work demonstrating synergism between these two drugs with regard to reinforcing effects [40] and discriminative stimulus effects [14,39]. Although it cannot be entirely ruled out based on the current studies, it is unlikely that self-administered caffeine, alone, was able to produce deficits in recognition memory given the body of work demonstrating either no change or enhancements in cognition rather than deficits following caffeine administration [58–70]. One potential mechanism underlying the ap-parent synergism between MDPV and caffeine to produce deficits in recognition memory is via enhancements in dopamine accumulation through the ability of caffeine to disinhibit dopamine D2 signaling. Perhaps this resulted in some degree of dopamine toxicity often associated with drugs acting as substrates for monoamine transporters (e.g., methamphetamine), wherein accumulated dopamine can form superoxide radicals and hydrogen peroxide that can result in neurodegeneration in regions associated with recognition memory behavior (e.g., hippocampus) (for a review, see [71]).

In an attempt to identify potential mechanisms underlying the behavioral changes, changes to striatal monoamine neurochemistry produced by each drug as a function of access condition were evaluated. With regard to the dopaminergic system, levels of striatal dopamine and DOPAC were largely unchanged independent of access condition or reinforcer available. There were small but significant reductions in HVA in rats that had 90-min access to cocaine, MDPV, or a mixture of MDPV+caffeine, whereas 90-min access to methamphetamine resulted in an increase in HVA, relative to rats with a history of 90-min access to grain pellets. 90-min access to methamphetamine also resulted in a significant increase in dopamine turnover. It should be noted that there was a significant negative correlation between methamphetamine intake and dopamine levels, however, no other such correlation was observed with regard to other reinforcers, or other neurotransmitters (data not shown), suggesting that it is not simply variability with regard to drug intake responsible for the lack of effects observed. Whereas dopamine neurochemistry was largely unchanged, all drugs appeared to reduce levels of striatal serotonin regardless of access condition, without large changes in 5-HIAA levels, although serotonin turnover appeared to increase in rats allowed 12-h access to MDPV+caffeine. Lastly, norepinephrine levels appeared to be greater in rats that had self-administered methamphetamine in both 90-min and 12-h access sessions, relative to those responding for grain pellets, whereas access to any other drug appeared to be without effect on norepinephrine levels. In contrast, 90-min access to all drugs and 12-h access to methamphetamine resulted in an apparent de-crease in epinephrine levels, accompanied by reductions in norepinephrine turnover. Generally, the seeming lack of large effects observed on striatal monoamine neurochemistry following thirty days of drug-taking, particularly in rats provided 12-h access, was unexpected. Methodological differences make comparisons with non-contingent “binge” models difficult. For instance, differences in duration of access per session (e.g., 90-min, 6-h, 12-h), total experimental duration, and time between the final self-administration session and tissue collection likely all contribute to the range of effects observed across studies. Indeed, the current studies utilized a 3-week drug-free period prior to tissue collection, to capture long-lasting persistent effects on neurochemistry. Previous work suggests that the duration of this period might preclude observing large changes in striatal neurochemistry that have been observed following a shorter drug-free period [72–75]. In addition to striatal monoamine neurochemistry, monoamine neurochemistry, neurodegeneration, and neuroinflammation will be quantified in other brain regions (hippocampus, prefrontal cortex) that are as-sociated with recognition memory, to allow for a deeper characterization of the persistent effects of self-administered synthetic cathinones and what could be driving the deficits in novel object recognition observed.

## 5. Conclusions

Despite the prevalence of MDPV and caffeine in “Bath Salts” preparations, relatively little is known regarding the persistent consequences of their use. The current studies found that rats allowed 90-min or 12-h access to MDPV+caffeine, or 12-h access to methamphetamine exhibited deficits in recognition memory, whereas rats allowed to respond for cocaine or MDPV did not. These findings are in line with what has been previously observed in rats allowed to self-administer methamphetamine and suggest that MDPV and caffeine interact not only to enhance abuse-related endpoints (e.g., discrimination, reinforcement, and reinstatement) but also adverse effects ranging from cognitive impairment to lethality. Generally, there were few changes to striatal monoamine neurochemistry, suggesting that monoamine levels recovered over the 3-week drug-free period, or physiological systems were able to compensate for drug effects. Future work will use extended access self-administration as an exposure paradigm to investigate the effects of chronic self-administered stimulants and stimulant mixtures on cardiovascular parameters and stimulant pharmacokinetics with the ultimate goal of developing novel treatments to prevent and/or rescue deficits produced by chronic stimulant use.

## Author Contributions

Conceptualization, RWSJ and GTC; methodology, RWSJ, KSM, GTC; formal analysis, RWSJ and GTC; investigation, RWSJ, KL, NW, and BL; reagents, AG and KCR; writing—original draft preparation, RWSJ and GTC; writing—review and editing, RWSJ, KSM, and GTC; All authors have read and agreed to the published version of the manuscript.

## Data Availability Statement

The original contributions presented in the study are included in the article/supplementary material, further inquiries can be directed to the corresponding author.

## Acknowledgments

The authors would like to thank Melson Mesmin, Rachel DeSantis, Torie Campos, Juan Morales, and Karen Jimenez for their technical assistance in the completion of these studies.

